# *Cochliobolus miyabeanus* species-specific identification using nonribosomal peptide synthetase (NRPS) and polyketide synthase (PKS)-encoding genes associated with virulence to rice

**DOI:** 10.1101/2025.02.19.638526

**Authors:** Nur Ain Izzati Mohd Zainudin, Nor Azizah Kusai, Nur Baiti Abd Murad, Dongliang Wu, Bradford J. Condon, B. Gillian Turgeon

**Affiliations:** Department of Biology, Faculty of Science, Universiti Putra Malaysia, 43400 Serdang, Selangor, Malaysia; Laboratory of Sustainable Agronomy and Crop Protection, Institute of Plantation Studies, Universiti Putra Malaysia, 43400 Serdang, Selangor, Malaysia; Section of Plant Pathology & Plant-Microbe Biology, School of Integrative Plant Science, Cornell University, Ithaca NY, USA

**Keywords:** *Bipolaris oryzae*, brown spot disease, virulence, rice (*Oryza sativa* L.)

## Abstract

*Cochliobolus miyabeanus* causes brown spot disease of rice. For necrotrophic *Cochliobolus* spp., some small molecule products of nonribosomal peptide synthetases (NRPS) and polyketide synthases (PKS) are known pathogenicity and/or virulence factors, often acting in a host-specific manner. Previous whole genome analyses across *Cochliobolus* species identified 11 NRPS- and 21 PKS*-*encoding genes in *C. miyabeanus*. First purpose of the current study was to examine the effect of deletion of ten of the discontinuously distributed genes (corresponding to JGI protein IDs 4446, 5802, 6546, 7015, 9064, 41693, 41753, 83551, 98843 and 107726) on virulence of *C*. *miyabeanus* to rice. A second purpose, based on results of the first, was to develop a PCR method to specifically identify *C. miyabeanus* and to distinguish it from other closely related Cochliobolus spp. We show that deletion of seven of the genes resulted in mutants that were reduced in virulence compared to the wild-type strain WK1C. Wild-type developed large dark brown necrotic lesions surrounded by chlorotic halos, while the mutants produced smaller, light brown spots with chlorosis. These results suggest that the products of these genes contribute to brown spot disease. In addition, we identified *PKS* and *NPS* SNPs that discriminate between *C. miyabeanus* and other species of *Cochliobolus*.

## Introduction

*Cochliobolus miyabeanus* (*Bipolaris oryzae*) is the causal agent of brown spot disease of rice, a worldwide problem. In 1942-43, the disease contributed to the Bengal rice famine that caused starvation of more than two million people (Devereux 2000). In addition to rice (*Oryza* species), the pathogen also causes severe leaf spot disease on American wildrice (*Zizania* species), wheat, maize, and switchgrass (Claudia et al. 2016; Strange and Scott 2005). In *Cochliobolus* spp., secondary metabolites that have been linked to virulence are often structurally diverse and complex small molecules known as nonribosomal peptides (NRPs) and polyketides (PKs) that are synthesized by large multifunctional nonribosomal peptide synthetase (NRPS) or polyketide synthase (PKS) enzymes which connect simple monomeric building blocks (amino acids and acyl-CoAs and variants, respectively) by a cascade of condensation reactions (Gallo et al. 2013; Sayari et al. 2019). In *C. miyabeanus*, other genes reported to be involved in pathogenicity/virulence or response to host defenses include hydrophobins, cutinases, cell wall degrading enzymes, enzymes related to reactive oxygen species scavenging, and detoxification systems (Castell-Miller et al. 2016).

In previous work, genome mining revealed that *C. miyabeanus* encodes 11 *NPS*s and 21 *PKS*s (Condon et al. 2013). In this study we report deletion of 10 of these genes, chosen based on their distinctive phylogenetic distribution (see Figs. S5, S6 in Condon et al., 2013), and found that seven were involved in virulence of *C. miyabeanus* to rice. These include the products of NRPS proteins 98843 and 107726 and PKS proteins 4446, 5802, 9064, 41753, and 83551 (JGI protein ID numbers). We also found that five of the corresponding nucleotide sequences could be used as diagnostic species-specific PCR assays to distinguish *C. miyabeanus* isolates from isolates of *Cochliobolus heterostrophus*, *Cochliobolus carbonum*, *Cochliobolus victoriae*, *Cochliobolus hawaiiensis*, *Cochliobolus sativus*, *Cochliobolus eragrostidis*, *Cochliobolus geniculatus* and *Cochliobolus lunatus* (Table 1, S1).

**Table 1.**
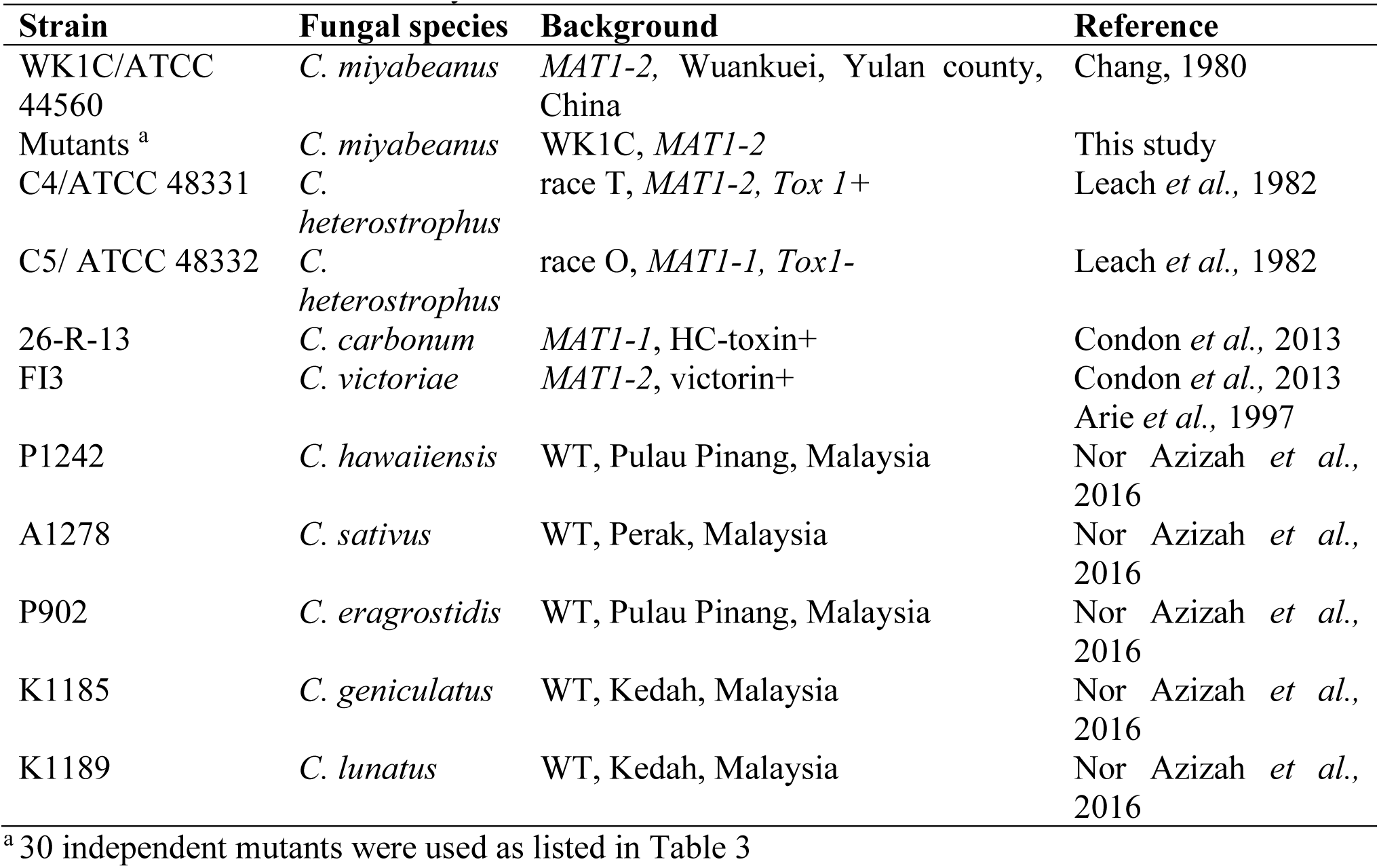
Strains used in this study.

## Material and Methods

### Fungal cultures and growth conditions

Fungal strains used in this study are listed in Table 1. All mutants were constructed in *C. miyabeanus* strain WK1C/ATCC44560 (Wuankuei, Yulan county, China, *MAT1-2*). For diagnostic PCR assays, DNA from nine additional *Cochliobolus* isolates was used: *C. heterostrophus* strains C4; ATCC 48331 (race T, *MAT1-2, Tox1+*), and C5; ATCC 48332 (race O, *MAT1-1, Tox1-*), *C. carbonum* strain 26-R-13 (race 1, *MAT1-1*, HC-toxin+), *C. victoriae* strain FI3 (unknown geographic origin, *MAT1-2*, victorin+), and five field strains, including *C. hawaiiensis* P1242, *C. sativus* A1278, *C. eragrostidis* P902, *C. geniculatus* K1185 and *C. lunatus* K1189 isolated from brown spot disease samples of rice in Peninsular Malaysia (Nor Azizah et al. 2016). All strains were revived from glycerol stock and were single conidiated on complete medium with xylose (CMX) (Tzeng 1990), and incubated at 28°C under 12/12 hour light/dark for 7 days.

### Selection of *C. miyabeanus* nonribosomal peptide synthetase and polyketide synthase encoding gene sequences for deletion

Ten individual PKS and NRPS-encoding genes of *C. miyabeanus* (PKS: JGI protein ID 4446, 5802, 6546, 7015, 9064, 41753, 83551; NRPS: 41693, 98843, 107726) were selected for deletion based on their distinctive placement in phylogenetic trees that included complete suites of these genes from 10 *Cochliobolus* strains (Table 2 and Condon et al. 2013, Figs. S5, 6). The candidate sequences were extracted from the *C. miyabeanus* WK1C genome sequence in mycocosm (https://genome.jgi.doe.gov/Cocmi1/Cocmi1.info.html) (Table 2).

**Table 2.** *C. miyabeanus NPS* and *PKS*-encoding gene sequences deleted.

| Node<sup>a</sup> | JGI Protein ID<sup>b</sup> | Type of protein | Scaffold and position | Relationship to *C. heterostrophus* strain C4 proteins<sup>c</sup> |
| --- | --- | --- | --- | --- |
| n155 | 7015 | NRPS:PKS | scaffold_107:80610-88238 | Pks24 |
| n281 | 41693 | NRPS | scaffold_313:598-12372 | Nps13amp 1, related to nps3 amp 2 |
| n311 | 41753 | PKS | scaffold_328:1201-8492 | unique |
| n1140 | 107726 | NPRS | scaffold_217:2744-6037 | LmSirP, Af GliP |
| n2370 | 83551 | PKS | scaffold_10:13147- 21235 | Pks6 |
| n2639 | 6546 | PKS | scaffold_94:147-1394 | Pks14 |
| n3591 | 9064 | PKS | scaffold_197:30760-36396 | Pks19/20 |
| n5640 | 5802 | PKS | scaffold_76:68169-69794 | ancestral to melanin Pks18 |
| n6511 | 4446 | PKS | scaffold_47:158475-165136 | Pks14 (ancestral to 2639) |
| n8391 | 98843 | NRPS | scaffold_91:54275-69751 | Nps3amps1-4 |

<sup>a</sup> Condon et al., 2013; nodes designated before JGI sequence was available.
<sup>b</sup> Joint Genome Institute (https://genome.jgi.doe.gov/Cocmil/Cocmil.home.html) protein ID corresponding to node (i.e., n155 = JGI protein ID 7015)
<sup>c</sup> Condon et al., 2013; Figs. S5, S6

### Fungal transformation

Transformation was carried out as previously described (Inderbitzin et al. 2010; Oide et al. 2006), with a slight modification for *C. miyabeanus*. WT and mutants strains were cultured on CMX for 14 days under constant light. For protoplasting, conidia were transferred to 300 ml complete medium (CM) in a 1L flask and shaken at 100 rpm for 20-24 hrs in the dark at 24°C. Protoplasts were harvested and transformed using the split marker approach as described previously (Catlett et al. 2003). Candidate transformants were single-conidiated and transferred to CMX without salts (CMNoS) containing hygromycin B (100 μg/ml) to confirm sensitivity to the drug, as described previously (Oide et al. 2006). All strains were stored in 25% CM glycerol at - 80°C.

### Gene deletions and confirmation of gene deletion

The split marker system (Catlett et al. 2003; Inderbitzin et al. 2010) was used for gene deletion. All primers were designed using Gene Runner 3.05 / 4.0.9.3 Beta software and Primer3 plus software (http://www.bioinformatics.nl/cgi-bin/primer3plus/primer3plus.cgi/). PCR amplification of transformation constructs was carried out with iProof high-fidelity DNA polymerase (Bio-Rad, Hercules, CA, U.S.A) following the manufacturer’s instructions.

To verify deletion of candidate genes, PCR amplifications of gene deletion were conducted with GoTaq polymerase (Promega, Madison, WI, USA) using primer pairs internal to each gene (Table S1). To further confirm integration of the hygromycin resistance gene at the target gene native locus, PCR was conducted using a primer located outside the 5’ and 3’ flanks used for gene deletion and second primer location in the hygromycin (*hygB)* cassette from pUCATPH (Lu et al. 1994), which confers resistance to hygromycin B (Catlett et al. 2003; Wu et al. 2012).

### Virulence assays

To evaluate the effect of deletion of each of the 10 genes on virulence to the host, five-week-old rice seedlings (variety Nipponbare) were inoculated by spraying plants with conidial suspensions at a concentration of 2 x 10^5^ conidia/ml in 0.02% Tween 20 (2 ml per plant). Four plants were used for each different fungal strain and pots were incubated in a mist chamber for 24 h after inoculation. The inoculated plants were then moved to a growth chamber for an additional 9 days at 24°C under a cycle of 16 h of light and 8 h of darkness. The inoculated leaves were photographed at 10 days after inoculation and the experiment was repeated three times. The data were analyzed using IBM SPSS analysis version 17.0 (IBM Coorporation, New York, United States).

### Development of diagnostic PCR assays for *C. miyabeanus*

We identified regions that had variable nucleotide sequences in the 10 genes we studied (7 *PKS* and 3 *NPS*) and designed PCR primers corresponding to five (protein IDs 5802, 4446, 83551, 9064 and 41753) of the 10 genes Table S1. These sequences are potentially species-specific.

PCR reactions using these primers and DNA from nine *Cochliobolus* spp. or isolates as template were run in thermal cycler (Biometra®Tprofessional, Göttingen, Germany) with the following cycle; 1 min for denaturation (93°C) and annealing, each (temperature depending upon primer), 2 min for synthesis (72°C) with longer denaturing step in the first cycle (3 min) with 7 min of synthesis in the last, and 30-35 cycles.

## Results

### Selection of *C. miyabeanus* nonribosomal peptide synthetase and polyketide synthase-encoding gene sequences for deletion

Ten individual PKS and NRPS-encoding genes of *C. miyabeanus* (PKS: JGI protein IDs 4446, 5802, 6546, 7015, 9064, 41753, 83551; NRPS: 41693, 98843, 107726) were selected for deletion based on their unique or discontinuous distribution in phylogenetic trees that included complete suites of PKS or NRPS proteins of 10 *Cochliobolus* spp. strains (Table 2, see Suppl. Figs. 5 and 6 in Condon et al. 2013). None of these is totally conserved across the *Cochliobolus* species or strains examined. The candidate sequences were extracted from the *C. miyabeanus* WK1C genome sequence in mycocosm (https://genome.jgi.doe.gov/Cocmi1/Cocmi1.info.html) as stated in Table 2.

### PCR verification of *NPS/PKS* gene deletion

Loss of complete or partial *NPS/PKS* genes and correct insertion of the selectable marker into the 5’ and 3’ regions flanking each target gene were verified by PCR using diagnostic primer pairs (Table S1, Fig. S1), as described in Materials and Methods and in Inderbitzin et al. (2010). For primer pairs internal to each gene, the expected PCR product was obtained from WT DNA, but not when DNA of the candidate deletion mutants was used as template. For confirmation of integration of the hygromycin resistance gene at the target gene native locus, no amplification occurred when WT DNA was used, while the expected amplicon was observed when DNA of the mutants was used as template (Fig. S1). Three independent mutants of each mutant (Δ107726 NRPS #1,#2, #4, Δ41693 NRPS #1, #3, #5; Δ41753 PKS #1, #3, #4; Δ4446 PKS #3, #5, #7; Δ5802 PKS #2, #4, #5; Δ6546 PKS #3, #5, #7; Δ7015 NRPS:PKS #3, #5, #7; Δ83551 PKS #1, #4, #5; Δ9064 PKS #1, #4, #5; Δ98843 NRPS #1, #4, #5) were used for subsequent evaluations.

### Seven of the *PKS/NPS* genes are required for full virulence of *C. miyabeanus* to rice

Lesion size and number were measured and characterized at 10-days post inoculation. Mutants deleted for seven out of 10 genes produced lesions that were statistically smaller and less numerous than those of WT (Table 3). These included mutants deleted for genes corresponding to PKS protein IDs 4446, 5802, 9064, 41753, and 83551 and NRPS IDs 98843, and 107726. Plants inoculated with mutants deleted for genes corresponding to the 6546 and 7015 PKS proteins and for the 41693 NRPS, produced lesions that were not statistically different from WT in terms of size, but were reduced in number. WT developed lesions with large dark brown cores surrounded by large beige necrotic areas and whose appearance was more necrotic overall compared to the mutants (Figs. 1, 2). For the mutants, lesion size and color varied depending on the isolate. Lesions produced by the mutant lacking PKS protein ID 41753 (independent strains #1, 3 and 4) were the smallest (0.914 - 1.066 ± 0.307 - 0.660 mm) with the fewest lesions (3.3 - 3.5 ± 1.252 - 1.509 per cm^2^). Other mutants, PKS-encoding genes corresponding to protein IDs 4446, 5802, 9064, 83551 (Fig. 1) and NRPS-encoding genes 98843 and 107726 (Fig. 2) produced significantly smaller and fewer lesions than those caused by WT (2.032 ± 0.905 mm, 12.6 ± 2.591 lesions per cm^2^). These findings suggest that these genes are producing metabolites necessary for normal virulence of *C. miyabeanus* on rice and thus, brown spot disease.

**Fig. 1.**
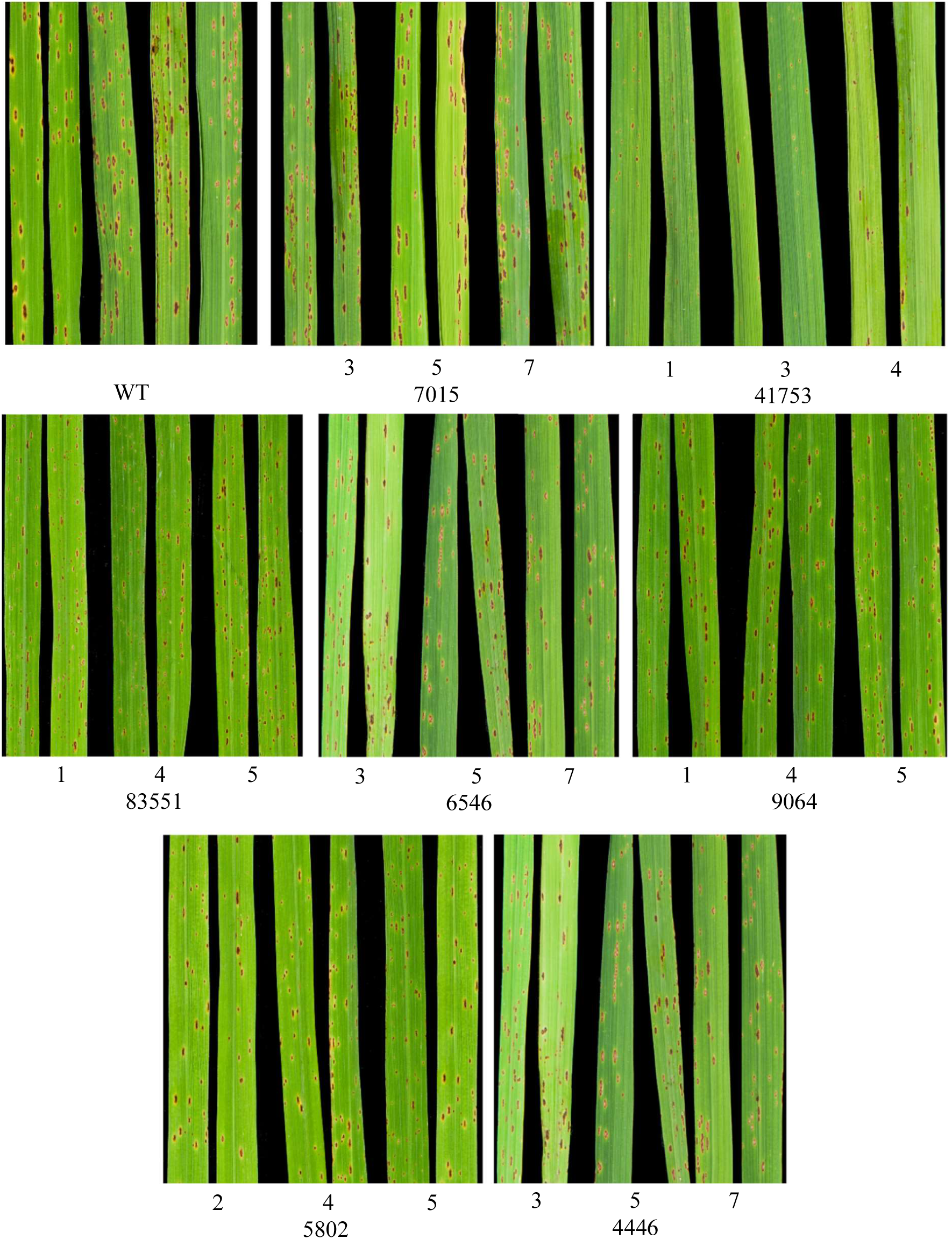
Virulence assays. Leaves inoculated with WT and *pks* mutants corresponding to deletion of genes corresponding to JGI protein IDs: 7015, 41753, 83551, 6546, 9064, 5802 and 4446. Δ41753, Δ4446, Δ5802, Δ83551, Δ9064 and Δ98843 mutants produced smaller and fewer lesions that were statistically different from WT. The leaves were photographed 10 days after inoculation.

**Fig. 2.**
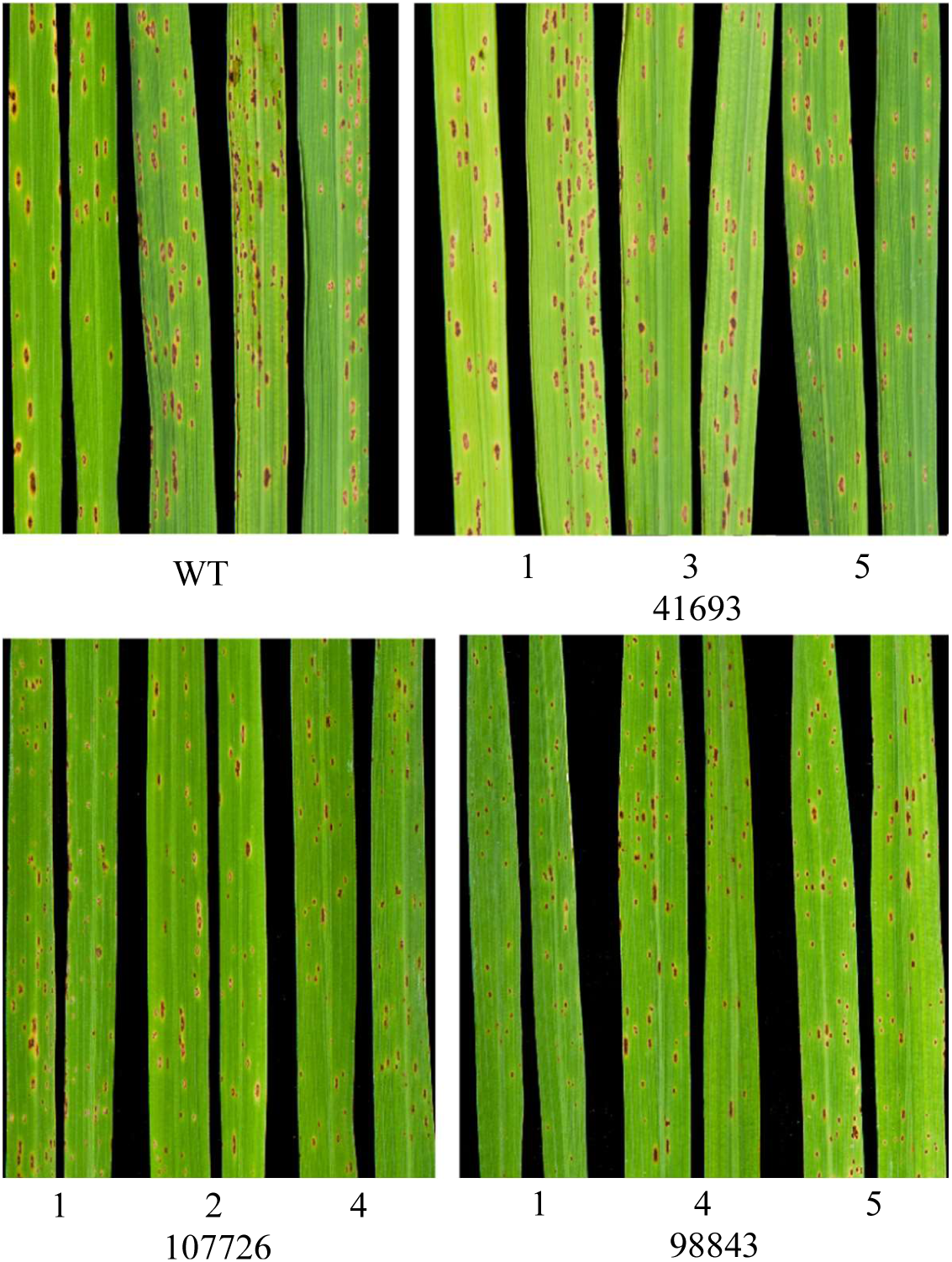
Leaves inoculated with WT and *nps* mutants corresponding to deletion of genes corresponding to JGI protein IDs 41693, 1007726 and 98843. Δ1007726 and Δ98843 mutants produced smaller and fewer lesions that were statistically different from WT. The leaves were photographed 10 days after inoculation.

**Table 3.**
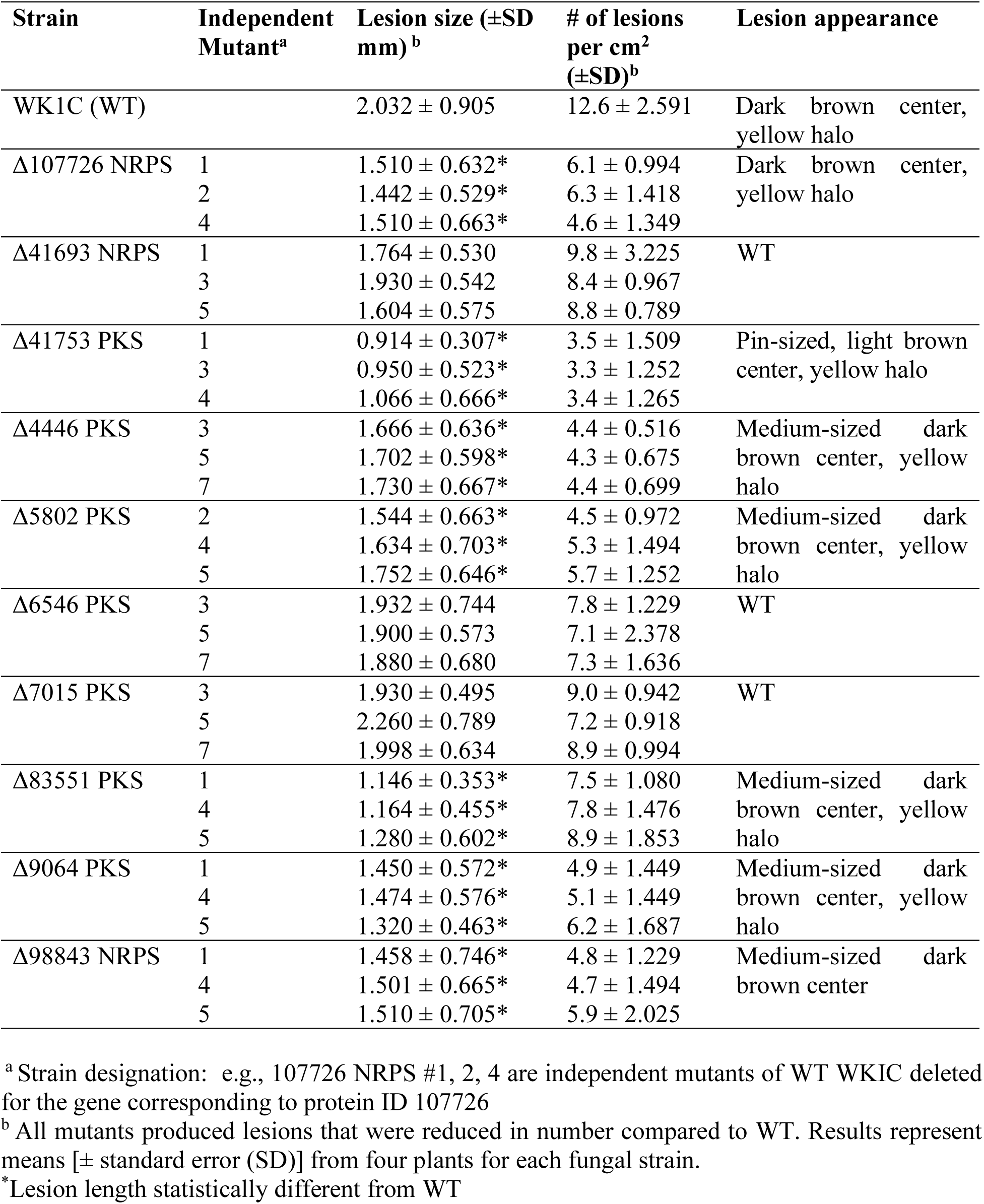
Virulence phenotype of *C. miyabeanus* mutants based on lesion length and number compared to WT.

### Five nucleotide sequences have SNPs suitable as diagnostic species-specific primers

Five pairs of primers were designed corresponding to positions that varied in nucleotide sequence between *C. miyabeanus* genes encoding protein IDs 4446, 5802, 9064, 41753, 83551 and other *Cochliobolus* species, to distinguish C. *miyabeanus* isolates from isolates of other species. When used with DNA templates from *C. miyabeanus*, *C. heterostrophus*, *C. carbonum*, *C. victoriae*, *C. hawaiiensis*, *C. sativus*, *C. eragrostidis*, *C. geniculatus* and *C. lunatus*, fragments were amplified only from *C. miyabeanus* (Table 4, Fig. 3). Thus, these five pairs of primers (NM132-NM133, NM71-NM72, NM57-NM58, NM114-NM115 and NM146-NM147, respectively) can be used for species-specific diagnostic PCR assays of *C. miyabeanus* (Table S1).

**Fig. 3.**
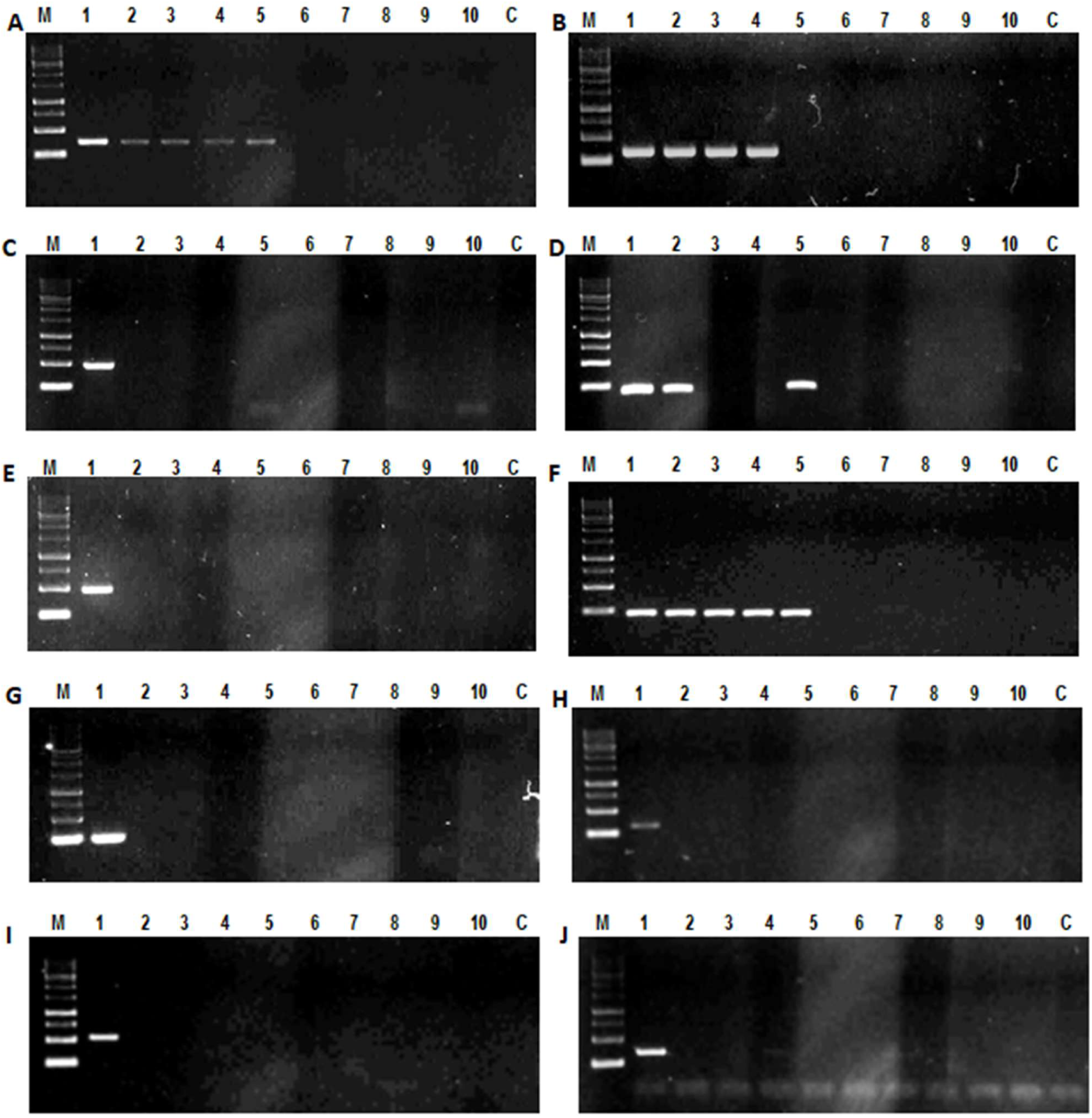
Diagnostic PCR amplification assay using DNA from several *Cochliobolus* species as template and species-specific primer pairs (Table S1). *nps/pks* mutants corresponding to JGI protein IDs, A= 7015, B=41693, C=41753, D=107726, E=83551, F=6546, G=9064, H=5802, I=4446, J=98843. Lane 1-13: M= Marker 1kb, 1=*C. miyabeanus* WK1C, 2=*C. victoriae* F13, 3=*C. heterostrophus* C4, 4=*C. heterostrophus* C5, 5=*C. carbonum* 26R13, 6=*C. hawaiiensis* P1242, 7=*C. sativus* A1278, 8=*C. eragrostidis* P902, 9=*C. geniculatus* K1185, 10=*C. lunatus* K1189; M= Marker 100bp.

**Table 4.**
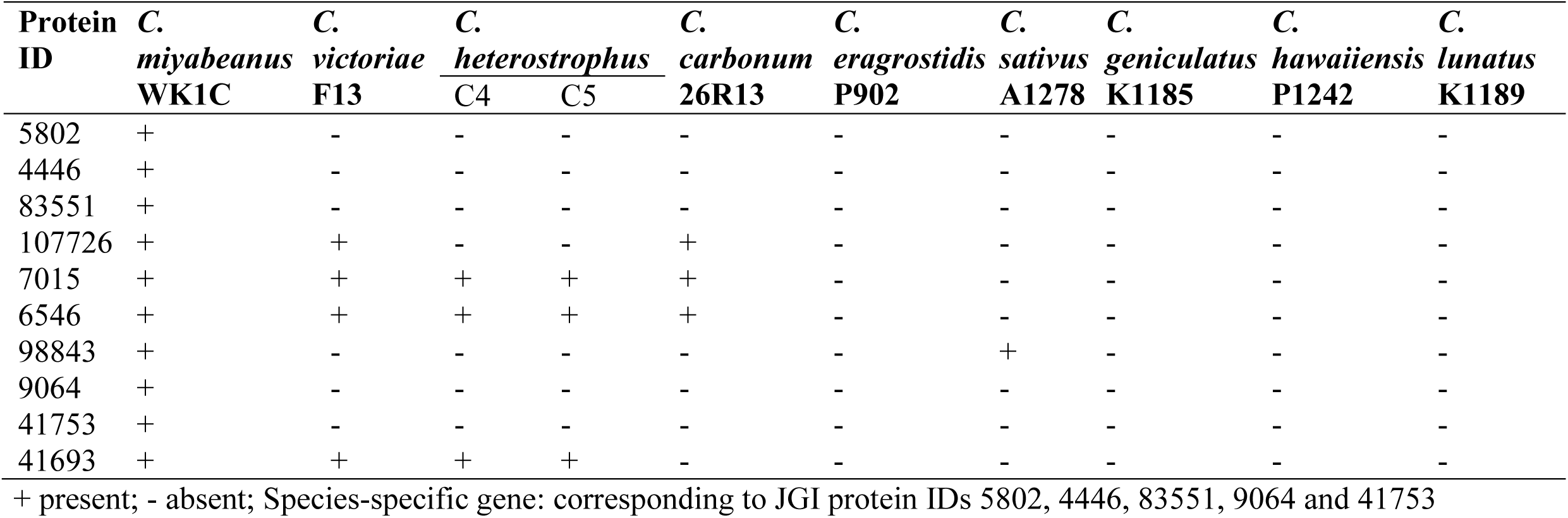
PCR assay with species-specific primers for *C. miyabeanus*.

## Discussion

Certain secondary metabolites are important for virulence of fungi to their hosts, including the polyketides associated with production of T toxin by *C. heterostrophus* race T (Haridas et al. 2023), the NRPS associated with production of HC-toxin in *C. carbonum* race 1 (Leonard and Leath 1990), and the NRPS produced extracellular siderophore NPS6 (Oide et al. 2006), to name a few. We explored whether or not mutants deleted for one of ten selected PKS/NRPS-encoding genes affected virulence to rice, i.e., do the metabolites produced by these genes contribute to brown spot disease? We found that deletion of genes corresponding to protein IDs 4446 (Pks14, ancestral to 2639), 5802 (outside melanin clade), 9064 (Pks19/20), 41753 (unique PKS), 83551 (Pks6), 98843 (Nps3) and 107726 (LmSirP, Af GliP) (Table 2, Condon et al. 2013 Figs. S5, 6) are required for normal virulence of the fungus to rice when compared to WT. Note that, like the *PKS*s associated with T-toxin, and the *NPS* associated with HC-toxin, the *C. miyabeanus* gene corresponding to protein ID 41753 is unique to that genome (not found in any other *Cochliobolus* species). This again argues that identifying unique SM genes is an effective strategy for identification of virulence determinants.

To our knowledge, this is the first report of PKS/NRPS-encoding genes being associated with brown spot disease caused by *C. miyabeanus,* in contrast to PKS/NRPS-encoding genes in other *Cochliobolus* species (Ahn et al. 2002; Condon et al. 2013, Lee et al. 2005; Oide et al. 2006). Further work is needed to elucidate the products of these enzymes and the mechanism by which they contribute to brown spot disease.

In the leaves inoculated with WT, large dark brown necrotic lesions surrounded by chlorotic halos developed, consistent with appearance of lesions previously observed in Zainudin et al. (2015). However, the mutants produced fewer, smaller, lighter brown, less chlorotic lesions compared to WT. Out of seven PKS/NRPS-encoding gene deletions, protein ID 41753_PKS mutants produced the smallest and fewest spots (Table 3). As noted above, this gene is unique to *C. miyabeanus* similar to the *PKS* genes associated with T-toxin, the *NPS* genes associated with HC-toxin, the *NPS* gene corresponding to protein 115356 involved in high virulence of *C. sativus* strain ND90Pr on barley cv. Bowman, and others, supporting the notion that identification of discontinuously distributed genes is a important strategy to identify virulence determinants.

Of the genes required for virulence, we identified five with variations between the nucleotide sequence in *C. miyabeanus* and other *Cochliobolus* spp. and designed species-specific PCR primers based on gene differences. Primer pairs were able to distinguish between *C. miyabeanus* and eight other species of *Cochliobolus* (*C. heterostrophus*, *C. carbonum*, *C. victoriae*, *C. hawaiiensis*, *C. sativus*, *C. eragrostidis*, *C. geniculatus,* and *C. lunatus)*. The five pairs of *C. miyabeanus*-specific primers (NM51-NM52, NM57-NM58, NM71-NM72, NM114-NM115 and NM132-NM133) amplified a fragment in *C. miyabeanus* WT, but did not amplify PCR products in the remaining *Cochliobolus* spp. considered. These *C. miyabeanus*-specific primers thus allow rapid and simple identification of *C. miyabeanus* isolates. Selection and breeding for resistance in cereals are facilitated by the use of species-specific markers to identify various fungi implicated in leaf spots, some of which are difficult to differentiate morphologically. All *Cochliobolus* species used in this study are associated with a wide range of diseases of diverse plant species (Ahn and Walton 1996; Arie et al. 1997; Baker et al. 2006; Chang 1980; Leach et al. 1982; Leng and Zhong 2012; Nor Azizah et al. 2016). The present *C. miyabeanus*-specific primers should be useful in this regard as they allow unambiguous and easy identification of *C. miyabeanus* isolated from the field.

## Acknowledgements

This work was partially supported by the Ministry of Higher Education under Fundamental Research Grant Scheme (grant no: FRGS/1/2012/STWN03/UPM/02/3/5524297). Nur Ain Izzati M. Z. was sponsored by UPM-MOHE postdoctoral fellowship while in the lab of BGT at Cornell University and Nor Azizah K. was sponsored by the graduate research fund GRF-UPM. The authors would like to thank the Joint Genome Institute for sequencing the genome of *C. miyabeanus*. Gene deletion and virulence assays were conducted while Nur Ain Izzati M.Z in Turgeon Laboratory at Cornell.

## DATA AVAILABILITY STATEMENT

The data that supports the findings of this study are available in the supplementary material of this article.

