## Supplemental Figure 1, Table 1 for "*Cochliobolus miyabeanus* species-specific identification using nonribosomal peptide synthetase (NRPS) and polyketide synthase (PKS)-encoding genes associated with virulence to rice"

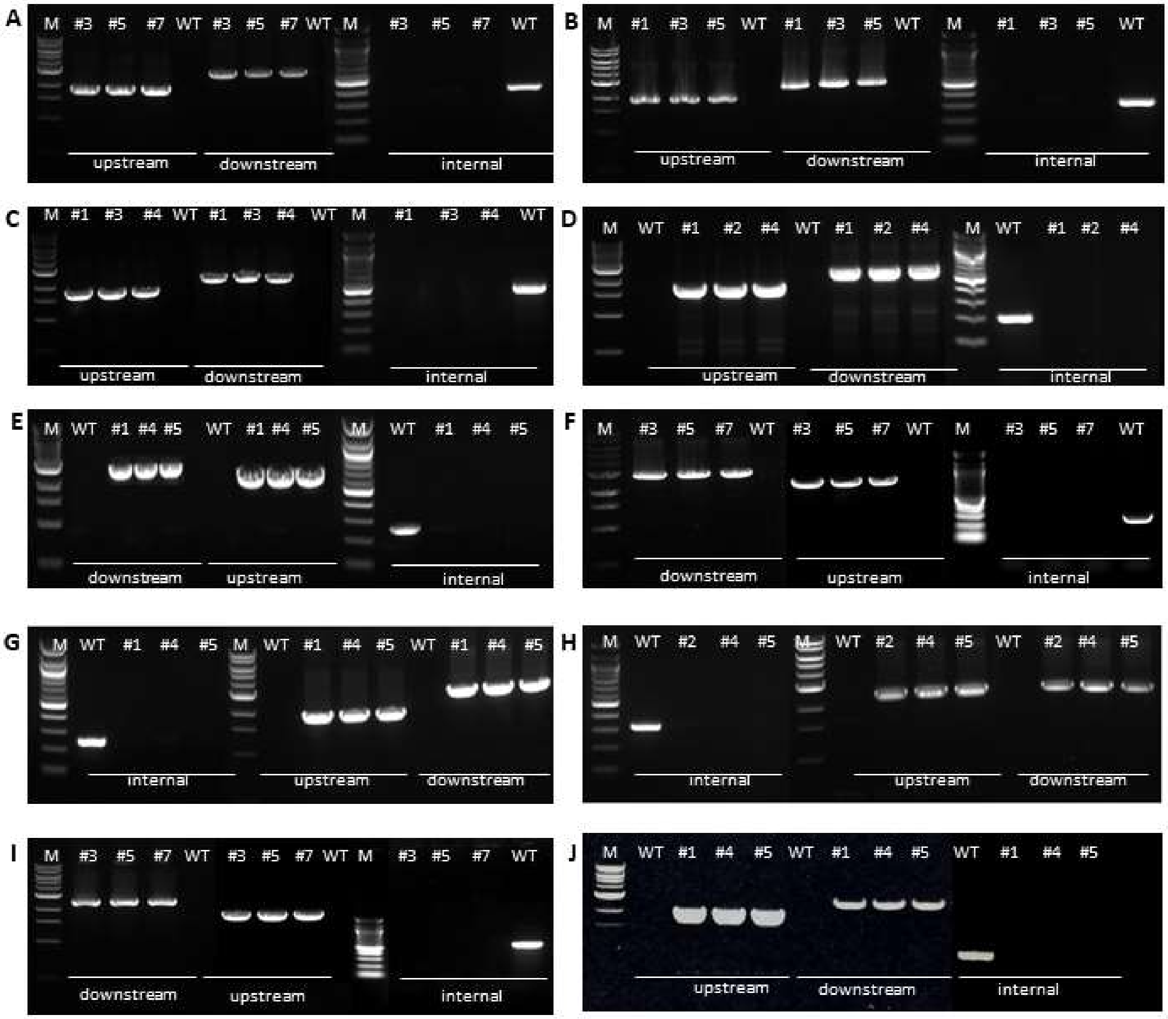


**Fig. S1**: Confirmation of gene deletion. Three pairs of primers (upstream, downstream, and gene-specific/internal) were used with genomic DNA of WT and three independent mutant strains as templates. A-J: NRPS/PKS nodes (protein ID), A = 7015, B = 41693, C = 41753, D = 107726, E = 83551, F = 6546, G = 9064, H = 5802, I = 4446, J = 98843. Assays confirmed deletions of all 10 genes in *C. miyabeanus*. M lane is 1kb marker for upstream and downstream pairs, or 100bp marker for specific internal fragment of the gene deleted.

**Table S1** Primers used in this study

| No. | Primer name | Primer sequence | Note | Expected fragment sizes |
|---|---|---|---|---|
| 1. | NM122 | ATGTCCGTTGGTTTGAGGT | Forward primer of 5’ flank for n155 7015 deletion | upstream fragment 657bp |
| 2. | NM123 | TCCTGTGTGAAATTGTTATCCGCTCACCTTGGACGATTTCAGA | Reverse primer of 5’ flank for n155 7015, complementary to M13R | |
| 3. | NM124 | GTCGTGACTGGGAAAACCCTGGCGAGGGATGCTCACCGAAAT | Forward primer of 3’ flank for n155 7015, complimentary to M13R | downstream fragment 745bp |
| 4. | NM125 | GCGGCAGAAAGTATCCAAG | Reverse primer of 3’ flank for n155 7015 | |
| 5. | NM126 | AGGATGCGTGGAAACAGAT | Forward primer for verification n155 7015 gene deletion | 366bp |
| 6. | NM127 | TTCCAGATACTTGCCCAGAC | Reverse primer for verification n155 7015 gene deletion | |
| 7. | NM116 | GTTTGGGGCTTAGGCAGTA | Forward primer of 5’ flank for n281 41693 deletion | upstream fragment 750bp |
| 8. | NM117 | TCCTGTGTGAAATTGTTATCCGCTCCTCTGTGTCGTCAAGTGC | Reverse primer of 5’ flank for n281 41693, complementary to M13R | |
| 9. | NM118 | GTCGTGACTGGGAAAACCCTGGCGGGTGCGGCATATCTGTTT | Forward primer of 3’ flank for n281 41693, complimentary to M13R | downstream fragment 470bp |
| 10. | NM119 | CTCGCTTATCCGTCACGTT | Reverse primer of 3’ flank for n281 41693 | |
| 11. | NM120 | GTAGGTGGCGAACTCATCC | Forward primer for verification n281 41693 gene deletion | 332bp |
| 12. | NM121 | CCTGTCCGCCCTATCTCTA | Reverse primer for verification n281 41693 gene deletion | |
| 13. | NM43 | AACATCCCATCACCACTCC | Forward primer of 5’ flank for n311 41753 deletion | upstream fragment 689bp |
| 14. | NM21 | TCCTGTGTGAAATTGTTATCCGCTTACTCAAAATCGTCGCCC | Reverse primer of 5’ flank for n311 41753, complementary to M13R | |
| 15. | NM22 | *GTCGTGACTGGGAAAACCCTGGCG*ACTCTGGGACTGGTGGTGT | Forward primer of 3’ flank for n311 41753, complimentary to M13R | downstream fragment 811bp |
| 16. | NM44 | AGCCGATTCACCATAGAGG | Reverse primer of 3’ flank for n311 41753 | |
| 17. | NM114 | CCTACACCCTAAGCGAACG | Forward primer for verification n311 41753 gene deletion | 479bp |
| 18. | NM115 | GATGATTGCCTCCTCTGCT | Reverse primer for verification n311 41753 gene deletion | |
| 19. | NM93 | GAACAACCACCAAGCCATT | Forward primer of 5’ flank for n1140 107726 deletion | |

| | | | | |
|---|---|---|---|---|
| 20. | NM94 | TCCTGTGTGAAATTGT TATCCGCTCACAATCT CCCCATAGCC | Reverse primer of 5' flank for n1140 107726, complementary to M13R | upstream fragment 691bp |
| 21. | NM95 | GTCGTGACTGGGAAA ACCCTGGCGAGCACTC GCCCTGTCTATG | Forward primer of 3' flank for n1140 g107726, complimentary to M13R | downstream fragment 669bp |
| 22. | NM96 | GCTCTATTGGACGGGT TTG | Reverse primer of 3' flank for n1140 107726 | |
| 23. | NM97 | CGTCTGGCATCAACTG TCT | Forward primer for verification n1140 g107726 gene deletion | 202bp |
| 24. | NM98 | AAGGCTGGTGAGGTTG TGT | Reverse primer for verification n1140 107726 gene deletion | |
| 25. | NM142 | GCCGCAGATACCAAG AGAG | Forward primer of 5' flank for n2370 83551 deletion | upstream fragment 752bp |
| 26. | NM143 | TCCTGTGTGAAATTGT TATCCGCTCCTTGCCT AACCTGTCGTC | Reverse primer of 5' flank for n2370 g83551, complementary to M13R | |
| 27. | NM144 | GTCGTGACTGGGAAA ACCCTGGCGGGTTATG GTCCCTTTGTCG | Forward primer of 3' flank for n2370 83551, complimentary to M13R | downstream fragment 787bp |
| 28. | NM145 | GGACCGTAAGTGGAG GAGA | Reverse primer of 3' flank for n2370 83551 | |
| 29. | NM146 | AGAGGCAAAACCGTG ACTT | Forward primer for verification n2370 83551 gene deletion | 490bp |
| 30. | NM147 | TGATCTCGGTGGTCTT TCC | Reverse primer for verification n2370 83551 gene deletion | |
| 31. | NM134 | GCAAAGAGTGGCAGG ATTC | Forward primer of 5' flank for n2639 6546 deletion | upstream fragment 953bp |
| 32. | NM135 | TCCTGTGTGAAATTGT TATCCGCTGTTGCCTT GCTCCTAATCC | Reverse primer of 5' flank for n2639 6546, complementary to M13R | |
| 33. | NM136 | GTCGTGACTGGGAAA ACCCTGGCGGGTTACG GAGGCACTAGAGG | Forward primer of 3' flank for n2639 6546 complimentary to M13R | downstream fragment 953bp |
| 34. | NM137 | TCTTGCCATCCGAGTT TG | Reverse primer of 3' flank for n2639 6546 | |
| 35. | NM138 | GGCCATGTTAATGGTG GTT | Forward primer for verification n2639 6546 gene deletion | 220bp |
| 36. | NM139 | TGGTTTCGTGATAGGC TGA | Reverse primer for verification n2639 6546 gene deletion | |
| 37. | NM45 | GCGACTCAGGTTCTAC GA | Forward primer of 5' flank for n3591 9064 deletion | upstream fragment 799bp |
| 38. | NM32 | *TCCTGTGTGAAATTGTT ATCCGCT*CGAGAACAG AATCCCACCT | Reverse primer of 5' flank for n3591 9064, complementary to M13R | |
| 39. | NM33 | *GTCGTGACTGGGAAAAC CCTGGCG*CTACGCTGA CAAATACCGC | Forward primer of 3' flank for n3591 9064 complimentary to M13R, | downstream fragment 750bp |

| | | | | |
|---|---|---|---|---|
| 40. | NM46 | AGCTTCTCTGCGATGACC | Reverse primer of 3' flank for n3591 9064 | |
| 41. | NM57 | GCGTCTTGCGTTAGTCAC | Forward primer for verification n3591 9064 gene deletion, | 249bp |
| 42. | NM58 | CATGCGGTGTCTATCGAG | Reverse primer for verification n3591 9064 gene deletion | |
| 43. | NM23 | GGTGTCGTCTTTTTGGTTTC | Forward primer of 5' flank for n5640 5802 deletion | upstream fragment 1210bp |
| 44. | NM24 | *TCCTGTGTGAAATTGTTATCCGCT*CGATGTGGATTGTGTCTGA | Reverse primer of 5' flank for n5640 5802, complementary to M13R | |
| 45. | NM25 | *GTCGTGACTGGGAAAACCCTGGCG*CGTCGTCATCATCATCATC | Forward primer of 3' flank for n5640 5802 complimentary to M13R, | downstream fragment 1241bp |
| 46. | NM26 | CTCCCAACAGCGTCTTCT | Reverse primer of 3' flank for n3591 9064 | |
| 47. | NM71 | CGCTTTTCATCTCCATCC | Forward primer for verification n5640 5802 gene deletion | l323bp |
| 48. | NM72 | CCACAGGCACAAAATGACA | Reverse primer for verification n5640 5802 gene deletion | |
| 49. | NM128 | GCTAGGCTCTCCAATCCAG | Forward primer of 5' flank for n6511 4446 deletion | upstream fragment 719bp |
| 50. | NM129 | TCCTGTGTGAAATTGTTATCCGCTCACACTTTGCTCTTCCCCTA | Reverse primer of 5' flank for n6511 4446, complementary to M13R | |
| 51. | NM130 | GTCGTGACTGGGAAAACCCTGGCGCATTAGCACCCACGACCTT | Forward primer of 3' flank for n6511 4446 complimentary to M13R | downstream fragment 705bp |
| 52. | NM131 | AAGTGAATCTTCCTGCCAGA | Reverse primer of 3' flank for n6511 4446 | |
| 53. | NM132 | ATCCCGTAGCGAATGTGTT | Forward primer for verification n6511 4446 gene deletion | 535bp |
| 54. | NM133 | CGACATCAGCAACAAGGAA | Reverse primer for verification n6511 4446 gene deletion | |
| 55. | NM75 | GCCGTTTTGCTGCTATCTT | Forward primer of 5' flank for n8391 98843 deletion | upstream fragment 890bp |
| 56. | NM76 | TCCTGTGTGAAATTGTTATCCGCTAAGGTGTCAGAACGCTACG | Reverse primer of 5'flank for n8391 98843 deletion, complementary to M13R | |
| 57. | NM77 | GTCGTGACTGGGAAAACCCTGGCGGGATGAGGACAAGGAAAGG | Forward primer of 3'flank for n8391 98843 deletion, complementary to M13R | downstream fragment 803bp |
| 58. | NM78 | GCGGGTGTGTGAATAAAGA | Reverse primer of 3' flank for n8391 98843 deletion | |
| 59. | NM83 | TCGTGTCAATCAGCAGAGA | Forward primer for verification n8391 98843 gene deletion | 360bp |
| 60. | NM84 | ACAGTGCCCAGTTGACATC | Reverse primer for verification n8391 98843 gene deletion | |
